# Effects of different pretreatments on flavonoids and antioxidant activity of *Dryopteris erythrosora* leave

**DOI:** 10.1101/353839

**Authors:** Xinxin Zhang, Xin Wang, Minglong Wang, Jianguo Cao, Jianbo Xiao, Quanxi Wang

## Abstract

Flavonoids with wide bioactivity for medcine are vital secondary metabolite of plant. The factors influenced on flavonoids had been reported. However, as the key processes lead to metabolite alterations, the influences of the different pretreatments of samples on flavonoids and antioxidant activity of ferns were with little information. Therefore, *Dryopteris erythrosora* leaves were chosen as the materials for analyzing flavonoids alterations, which would not only provide the significant basic data for flavonoid metabolism of fern, but also for further developing fern resources. The results showed that a) The total flavonoids contents of *D. erythrosora* leaves with different pretreatments were obviously different. The total flavonoid contents of samples, which was dried in shade firstly and then dried at 75 °C in oven, finally smashed, was the highest (7.6%), but that of samples, which was quickly dried at 75 °C in oven directly after cleaning and then smashed, was the lowest (2.17%); b) Antioxidant activities of *D. erythrosora* leaves with different pretreatments were variant. Samples, which were dried in shade firstly and then dried at 75 °C in oven, finally smashed and samples which were firstly dried in the sun and then dried at 75 °C in oven, ultimately smashed, both showed stronger antioxidant activity; c) Total twenty-three flavonoids with four different pretreatments were tentatively identified by HPLC-ESI-TOF-MS. In conlusion, a) The influences of different pretreatments on flavonoids and antioxidant activity of *D. erythrosora* Leaves were obvious. b) The best pretreatment in respect to conserving fern medical application was drying in shade firstly and then drying at 75 °C in oven, finally smashed.

## Introduction

Flavonoids are the important secondary metabolites of plant and extensive bioactivities for medicine [1], and would be influenced by kinds of factors, such as harvesting times, ecology factors [2–4], shade nettings and sowing time [5], development [6], light transmittance paper bags [7]. Besides, sample pretreatment was the key process leading to alterations in the quantity and quality of bioactive compounds [8–10]. Flavonoid content of fresh mulberry leaves was highest and that of leaves with oven-dried at 100-105 °C lowest [11]. Flavonoid contents, DPPH scaveging activities and reducing power of *Salvia officinalis* L. drying at shade were higher than that of samples drying in oven at 65 °C [12]. *Paramignya trimera* drying oven at 25 °C showed higher flavonoid contents than samplas drying in a heating oven at 100 °C [8]. Flavonoid yields of *Belamcanda chinensis* with different temperatures ranging from 40 °C to 120 °C in 10 °C intervals using a thermostatic oven were different [13]. Above reports obviously showed different pretreatments lead to alterations in flavonoid content and biological activity of plants.

However, how did the different pretreatments affect the flavonoid and bioactivities variation was still unclear. In addition, ferns, as the representative of plants with high flavonoid contents, was with little information on the effects of different pretreatments on flavonoids. Therefore, *Dryopteris erythrosora* (Eaton) O. Ktze. leaves were chosen as materials for the analysis of flavonoids. The aims of this study were to (I) assess the effects of different pretreatments on flavonoids and antioxidant activity of *D. erythrosora* leaves and (II) determine the best pretreatment in respect to conserving fern medical application, which would not only provide significant basis for the metabolite of fern flavonoids, but also further developing fern resources.

## Materials and methods

### Plant materials

*D. Erythrosora* leave in half-shade habitat were collected from Shanghai Sheshan National Forest Park in April 2017. The plants were identified by Prof. Jianguo Cao. Voucher specimens were deposited in the STC of the College of Life & Environmental Science, Shanghai Normal University.

### Chemicals

The chemicals were the same with previous report [14]. Rutin (purity > 99.0%), 2,2-Diphenyl-1-picrylhydrazyl (DPPH), 2,2’-azinobis-(3-ethylbenzothiazoline-6-sulfonic acid) (ABTS), Nitrotetrazolium blue chloride (NBT), phenazine methosulfate (PMS), nicotinamide adenine dinucleotide (NADH), 5, 5’-dithiobis-(2-nitrobenzoic acid) (DTNB) and 2,4,6-tri-2-pyridyl-s-triazine (TPTZ) were purchased from Sigma Co. (Shanghai, China). Acetonitrile was purchased from Thermo Fisher Scientific (Shanghai, China).

### Preparation of plant extracts

Fresh *D. Erythrosora* leaves were seperated into four groups at random and then treated with four ways respectively. Samples of group A were quickly smashed directly with liquid nitrogen after cleaning. Samples of group B were dried in shade firstly, then dried at 75 °C in oven for 48 h, finally smashed. Samples of group C were firstly dried in the sun (some old newspapers were flattened on the open area and then the samples were spread on the old newspapers beneath the sun), then dried at 75 °C in oven for 48 h, ultimately smashed. Samples of group D were quickly dried at 75 °C in oven for 48 h directly after cleaning, then smashed. Smashed samples were filtered by a 80-mesh screen.

1.00 g powders weighed from group B, group C and group D, respectively, were added with 60% ethanol (25 mL) for extraction at 50 °C for 2h with the ultrasound-assisted (20 min), respectively. the extraction processes were repeated twice, the mixture was filtered via a vacuum suction filter pump and the volume of the solution kept constant at 50 mL. In order to getting same dry weight, 3.3 g samples from group A were pounded in a mortar after calculating the drying rate, and then were added with 60% ethanol (25 mL) for extraction, the operation was the same as above processes.

One part of the extracts was used for determining the total flavonoid content and antioxidant activities, the other was extracted respectively by petroleum ether (PE), dichloromethane (DCM), ethyl acetate (EtOAc) and n-butanol (nBuOH) for the HPLC-ESI-TOF-MS analysis of flavonoids.

### Determination of total flavonoids content

A colorimetric assay was adopted for the determination of flavonoids content. Firstly, gradient concentration rutins were successively added 5% NaNO_2_ for 6 min, 5% Al(NO_3_)_3_ for 6 min, 4% NaOH for 12 min, and then the optical density (OD) of the mixture were recorded at 510 nm. Secondly, the linear equation (y=A+Bx) of rutin was plotted with the software Origin 7.5. Thirdly, the optical density (OD) of the gradient concentration extracts were determined. At last, accordting to the following formula, the total flavonoid content were calculated [14]. The formula was used as follows: total flavonoid content (%) = [(OD_1_+OD_2_+OD_3_)/3-A]/B*10/2*Volume/1000*100%

## Antioxidant Activity

### DPPH Assay

The DPPH free radical scavenging activity assay refered to previous report [14]. Briefly, 1 mL DPPH (0.1 mM in ethanol) and extracts with gradient concentration mixed firstly, the absorbance value was measured at 517 nm after incubating for 30 min. 60% methanol substituted for sample were taken as the control group. The DPPH free radical scavenging activity was calculated using the following formula: DPPH free radical scavening activity (%) = (1-A_sample 517_/A_control 517_)*100. The experiments were performed in triplicate with similar results (RSD < 5.0%).

### ABTS Assay

The ABTS assay of the extracts also refered to previous report [14]. In Short, 150 μL extracts with gradient concentration and 3mL appropriately diluted ABTS solutions mixed. The absorbance value at 734 nm was determined after incubating for 6 min. The ABTS free radical scavenging activity was calculated using the following formula: ABTS free radical scavening activity (%) = (1-A_sample 734_/A_control 734_)*100. The experiments were performed in triplicate with similar results (RSD < 5.0%).

### Superoxide Anion (O^2-^) Scavenging Activity

The determination of superoxide anion (O^2-^) scavenging activity was the same with previous report [14]. In brief, 1mL extracts with gradient concentrition mixed with sodium phosphate buffer were added with 1mL NBT (150 μM), 1mL NADH (468 μM) and 1mL PMS (60 μM) in turns, incubating at 25 °C for 5 min. The absorbance at 560 nm was determined. Superoxide anion (O^2-^) scavenging activity was calculated using the following formula: Superoxide anion (O^2-^) scavenging activity (%) = (1-A_sample 560_/A_control 560_)*100. The experiments were performed in triplicate with similar results (RSD < 5.0%).

### Reducing Power Assay

The reducing power assay was the same with reported method previously [14]. Briefly, The mixture including 1mL extract with gradient concentration, 2.5 mL phosphate buffer and 2.5 mLpotassium ferricyanide was put in water bath at 50 °C for 20 min, then 10% TCA was added to terminate the reaction. After centrifugation, one half of supernatant was mixed with 2.5 mL of distilled water and 0.5 mL of 0.1% ferric chloride, but the other half of supernatant were added with 3 mL of distilled water as the control group. Optical density at 700 nm reflected the reducing power. The experiments were performed in triplicate with similar results (RSD < 5.0%).

### FRAP Assay

The FRAP assay was also similar with our previous report [14]. The FRAP reagent was made up with TPTZ (10 mM) in HCl solution (40 mM) and FeCl_3_ (20 mM) in 250mL acetate buffer (pH 3.6). FRAP reagent should be used immediately after it was ready. Extracts of gradient concentration were added with FRAP reagent, respectively. After 4 mins, the optical density of the mixture at 593 nm was determined. With the calibration curves of Fe^2+^, the results were calculated. The FRAP reagent with distilled water was taken as the control group. The experiments were performed in triplicate with similar results (RSD < 5.0%).

### Flavonoids analysis of *D. erythrosora* leaves with different pretreatments by HPLC-ESI-TOF-MS

The Chromatographic separation was performedonan Agilent 1100 HPLC system (Agilent Technologies), equipped with abinary pump, amicrodegasser, Hi-performance well-plateauto sampler, thermostated column compartment and diode-array detector (DAD). UV spectra were recorded between 190 and 400 nm, and the UV detector was set at 254nm. Separation was performed on a SHISEIDO MG-C18 (1003.3 mm; i.d.3.0 mm) column using a gradient elution [methanol (A)/ water (0.1%HCOOH)(B)].

Extractions of petroleum ether (PE), dichloromethane (DCM), ethyl acetate (EtOAc) and n-butanol (nBuOH) were diluted 10 times respectively. The gradient program was 0-15 min,15-45% A;15-25 min, 45-55% A; 25-35 min, 55-90% A; the flow rate was kept at 0.4mL/min, and the sample injection volume was 10 μL and the column temperature was set at 25 °C. All MS experiments were conducted on an Agilent 6220 Time-of-Flight massspectrometry (TOF) equipped with an electrospray ionization (ESI) interface (Agilent Technologies, USA). Both the auxiliary and nebulizer gases were nitrogen with a flow rate of 10 L/min. The MS analysis was performed in both positive and negative scan modes under the following operation parameters: the dry gas temperature was set at 350 °C, voltage was 160V and the nebulizer pressure was set at 45 psi. Full scan data acquisition and dependent scan event data acquisition were performed from m/z 100-1200.

## Results and discussion

### Total flavonoid contents of *D. erythrosora* leaves with different pretreatments

The total flavonoid contents of *D. erythrosora* leaves with different pretreatments were determined as 7.38%, 7.6%, 6.41% and 2.17%, respectively (Fig 1), showing that the total flavonoid contents of extracts from group B which were dried in shade firstly, then dried at 75 °C in oven was the highest, but that of extracts from group D which were quickly dried at 75 °C in oven directly after cleaning was the lowest.

In the process of drying in shade, the rate of water loss was slow, the life process could not completely ended, so the plant could complete the flavonoid metabolite resulting in the increasing of total flavonoid content. However, in the process of drying in the sun, the rate of water loss was faster, the life process and flavonoid metabolite completely ended in short time, so the total flavonoid content did not increase. Full sunlight irradiance would influence flavonoid metabolite in leaves [15–16], which could result in the decrease of total flavonoid content. Samples from group A, which quickly smashed directly with liquid nitrogen after cleaning, showed the total flavonoid content of the live samples. It was speculated that water and sunlight might be main factors on total flavonoid content.

In addtion, flavonoids were found to be heat sensitive [17]. Heating at 75 °C directly could destroy enzyme activity and caused the synthesis pathway of flavonoids blocked, thus the total flavonoid content of Group D was the lowest.

### Antioxidant Activity

The DPPH free radical scavenging activities of the extracts of *D. erythrosora* leaves with different pretreatments were shown in Fig 2. When the doses were from 0 to 80μL, the DPPH free radical scavenging potential observably increased. Group C was the higher than group B. Less than 10 μL of the extracts could scavenge about 50% of the free radicals. However, group D was far lower than that of group C. The DPPH free radical scavenging activity of the extracts were arranged as group C>group B>group A>group D.

The ABTS free radical scavenging activities were shown in Fig 3. When the volume was between 0 and 150 μL, the ABTS free radical scavenging potential observably increased. Group B was obviously the highest and group D the lowest. 120 μL of the extract could scavenge about 50% of the free radicals. The ABTS free radical scavenging potential were arranged as group B>group C>group A>group D.

The superoxide anion scavenging activity were shown in Fig 4. With the range from 10 to 90 μL, the superoxide anion scavenging potential observably increased. The scavenging activity of four groups were similar, but group C was obviously higher than group A and group B, which showed samples which firstly dried in the sun and then dried at 75 °C in oven possessed stronger superoxide scavenging activity than others.

The results of FRAP assay and reducing power assay were shown in Figs 5 and 6. The results illustrated that the extracts from *D. erythrosora* leaves with different pretreatments both possessed antioxidant and reductive activity of Fe^3+^. With the volume increasing, the activity got improved. The activities level was arranged as group B>group C>group A>group D.

Above results illustrated that antioxidant activities of *D. erythrosora* leaves with different pretreatments were different. Except for superoxide anion (O^2-^) scavenging assay, group B and group C both showed stronger antioxidant activity than group D, which might be relative to the structure of flavonoids.

### Flavonoids analysis of *D. erythrosora* leaves with different pretreatments by HPLC-ESI-TOF-MS

HPLC-ESI-TOF-MS was used for the qualitative analysis of flavonoids of *D. erythrosora* leaves. By comparison with the known chromatograms and mass spectral data, a total of twenty-three peaks were tentatively identified as flavonoids and the detailed MS information were listed in Table 1, which contained eight flavonols, four flavones, three chalcones, two flavanols, two flavanones, two homoisoflavones, one isoflavone, one isoflavanone in the mass spectrometry-total ions chromatogram (MS-TIC) of extracts from *D. erythrosora* leaves with different pretreatments in negative ion mode. The results showed that the main flavonoids of *D. erythrosora* leaves were flavone and flavonols, which were consistent with the previous reports [18–22].

Chalcones and isoflavones, absent in group A, both be found in group B and group C, which was the first reported in Dryopteridaceae. This results showed chalcones and isoflavones could not be synthetised during *D. erythrosora* leaves natural growth, but could come out under stresses from sun drying or shade drying. On the side, except anthocyanin and its derivatives, all flavonoid types in flavonoid synthesis pathway could be found [23]. It was concluded that the flavonoids metabolic pathway of ferns was similar with spermatophyte, and metabolic pathway was closely relative to stress response.

Fourteen flavonoids from group A (Fig 7), twenty-three flavonoids from group B (Fig 8), fifteen flavonoids from group C (Fig 9) and six flavonoids from group D (Fig 10) were tentatively identified respectively. The results indicated that the loss of flavonoids from group D was greatest.

From Fig 7 to Fig 10, it was found that scutellarein 7-O-glucobioside, apigenin 7-O-rutinoside, apigenin-C-pentoside and kaempferol 3-O-α-L-arabinopyranoside were common flavonoids. That was to say, they were all unchanged with different pretreatments, which meant the flavonoid biosynthesis of four components were unaffected by the pretreatments. However, the contents of four flavonoids changed obviously (Fig 11). Taking samples quickly smashed directly with liquid nitrogen after cleaning as the standard, it was found that the content of scutellarein 7-O-glucobioside would increased, and the largest amount of which were obtained from samples which quickly dried at 75 °C in oven directly after cleaning. The results showed that the pertreatments of group B, group C and group D resulted in the synthesis of scutellarein 7-O-glucobioside and high temperature accelerated the synthesis. Scutellarein 7-O-glucobioside might be the product against high temperature. Meanwhile, it was observed that the contents of apigenin 7-O-rutinoside and kaempferol 3-O-α-L-arabinopyranoside both decreased, and the lowest content was in samples which quickly dried at 75 °C in oven directly after cleaning. This demonstated these pretreatments would made apigenin 7-O-rutinoside and kaempferol 3-O-α-L-arabinopyranoside decomposed or transformed. In addition, the amount of apigenin-C-pentoside increased in samples with the pretreatments of group B and group C but decreased in samples from group D, which meant the temperature might be the main factor influencing the synthesis of apigenin-C-pentoside.

Furthermore, If group D was neglected, seven common flavonoids, which did not contain four common flavonoids in four groups, would be found in group A, group B and group C. The contents of these flavonoids changed also obviously (Fig 12). Taking samples quickly smashed directly with liquid nitrogen after cleaning as the standard, the contents of myricetin 3-O-glucoside, kaempferide 3-rhamnoside-7-(6″-succinylglucose) and isocarthamidin-7-O-glucuronide increased, but the amounts of quercetin-3-O-β-D-xylopyranoside, apigenin-6-C-ara-8-C-glu reduced, which indicated that water might be the major influence factor. The content of koreanoside B changed little, which manifested that the synthesis of koreanoside B could not be enfluenced. The amount of rutin significantly decreased in group C, which stated that the light might result in the decomposition or the transformation of rutin.

Most notably, according to Table 1, it was found that the number of flavonoids were the greatest in group B. Comparing with group A, group B added nine flavonoids types and about the contents of 73% common compounds increased, which illustrated that the pretreatment of group B could not only result in the synthesis of other flavonoid types, but contribute to accumulate the content of flavonoids. Thereinto, especially, the content of kaempferide 3-rhamnoside-7-(6″-succinylglucose) was twice more than group A, which indicated that the synthesis of kaempferide 3-rhamnoside-7-(6″-succinylglucose) was the stress response of resistance to the pretreatment of group B.

The effects of high light irradiance on the biosynthesis of dihydroxy B-ring-substituted flavonoid had been reported [24–26]. Herein, comparing with group A, samples with the pretreatment of group B lost three flavonoids, but added quercetin-O-dihexoside, biochanin A-7-O-glucoside-6”-O-malonate, isoliquiritin apioside, 2S-5,7,2’,5’-Tetrahydroxy-6-methoxyflavanone, which meant light irradiance was indeed the sensitive factor to the biosynthesis of flavonoid. In different pretreaments, the loss of flavonoids in samples with pretreatment of group D were the greatest, 67% flavonoids lost.

## Conclusion

This paper explored the influences of different pretreatments on the total flavonoid contents, antioxidant activity and flavonoid ingredients of *D. erythrosora* leaves. The main conclusions were as follows: a) The total flavonoids contents, antioxidant activities and flavonoid ingredients of *D. erythrosora* leaves with different pretreatments varied obviously, and samples which dried in shade firstly, then dried at 75 °C in oven showed the highest flavonoid content and strongest antioxidant activities. b) The pretreatment which quickly dried at 75 °C in oven directly after cleaning would make great loss of flavonoids, and the best pretreatment in respect to conserving fern medical flavonoids application was drying in shade firstly, then drying at 75 °C in oven and finally smashed and filtered. This paper was vitally significant in guiding the development and utilizing of ferns.

## Acknowledgments

The work was sponsored by Shanghai Engineering Research Center of Plant Germplasm Resources (No. 17DZ2252700)

---

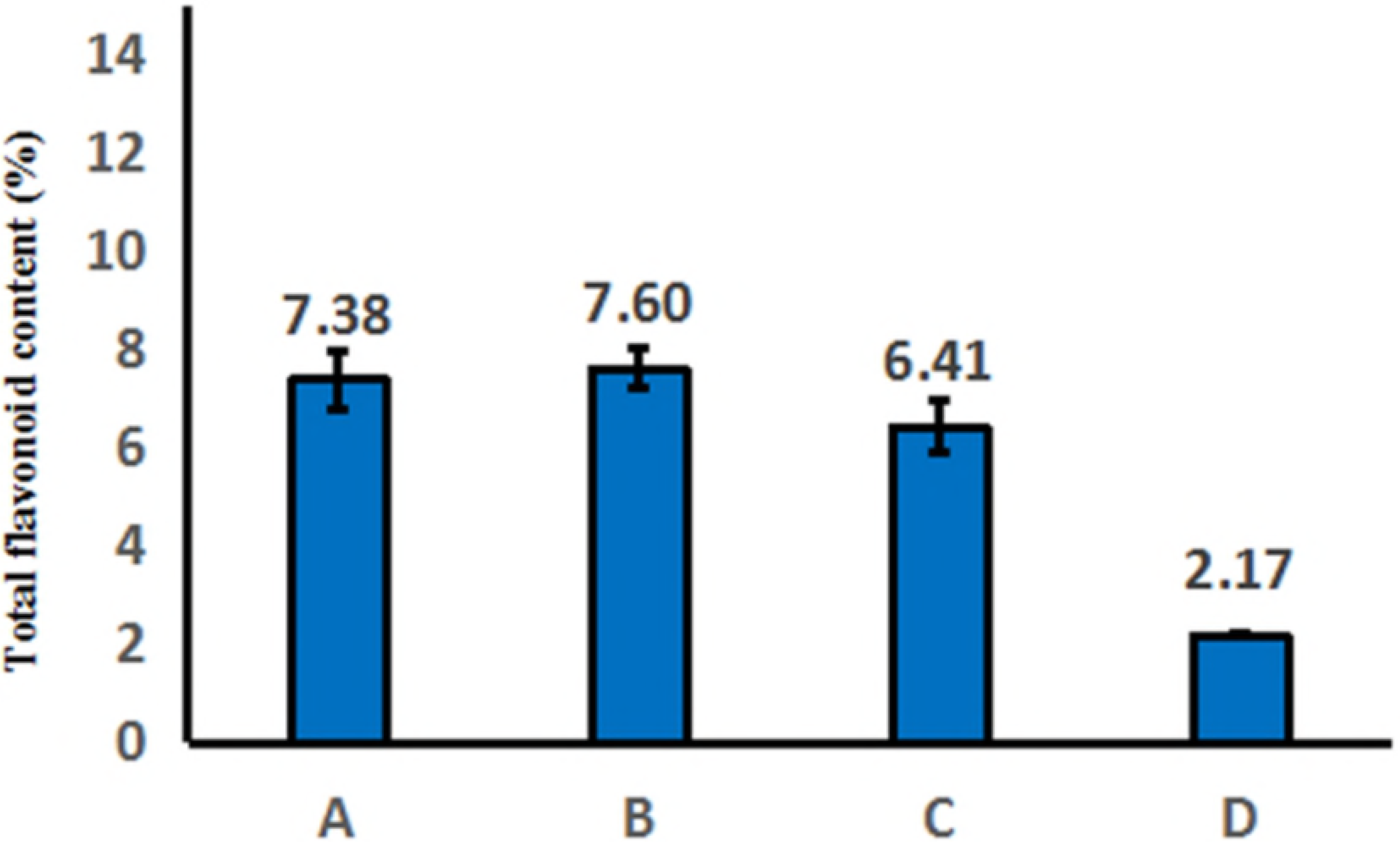

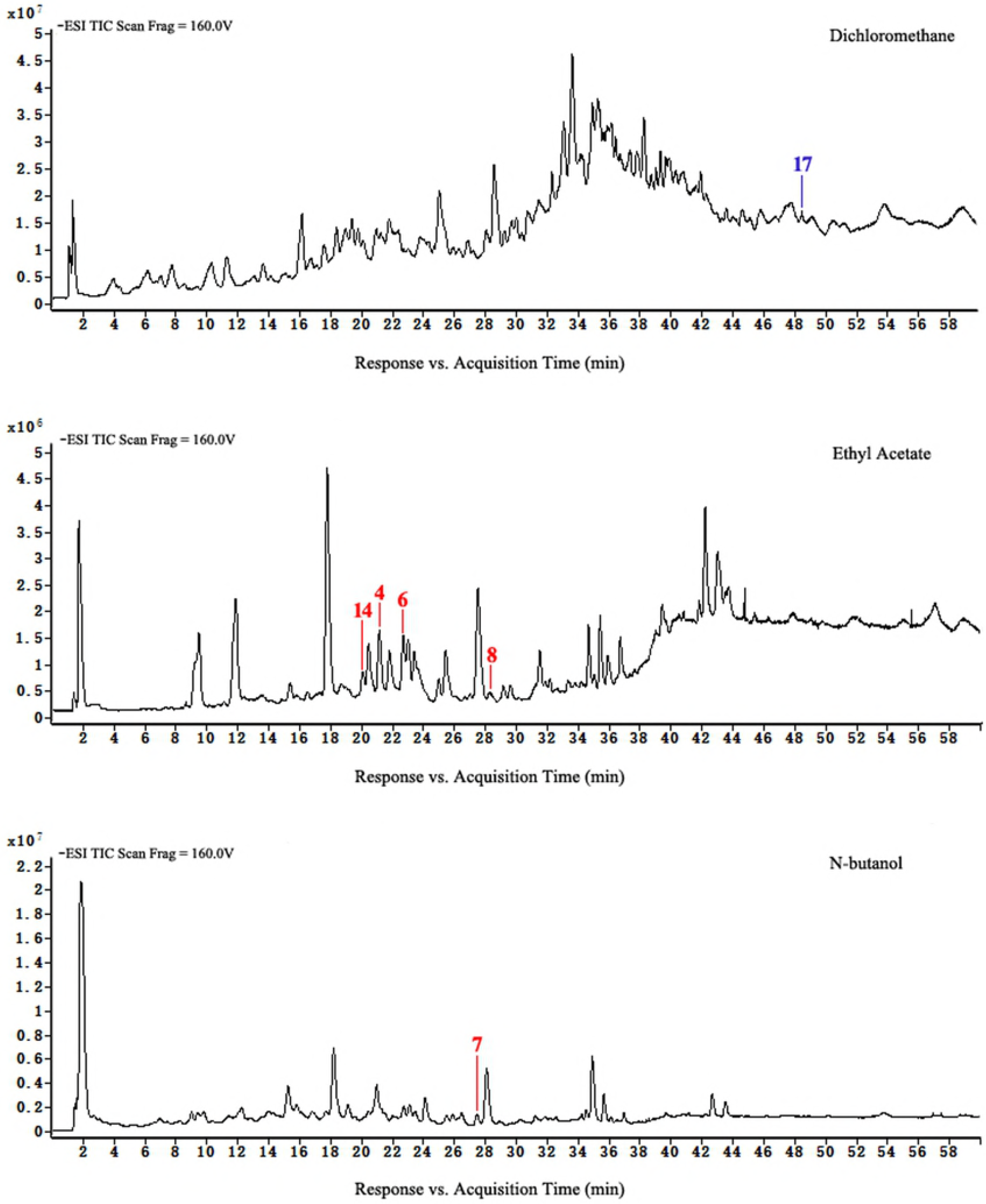

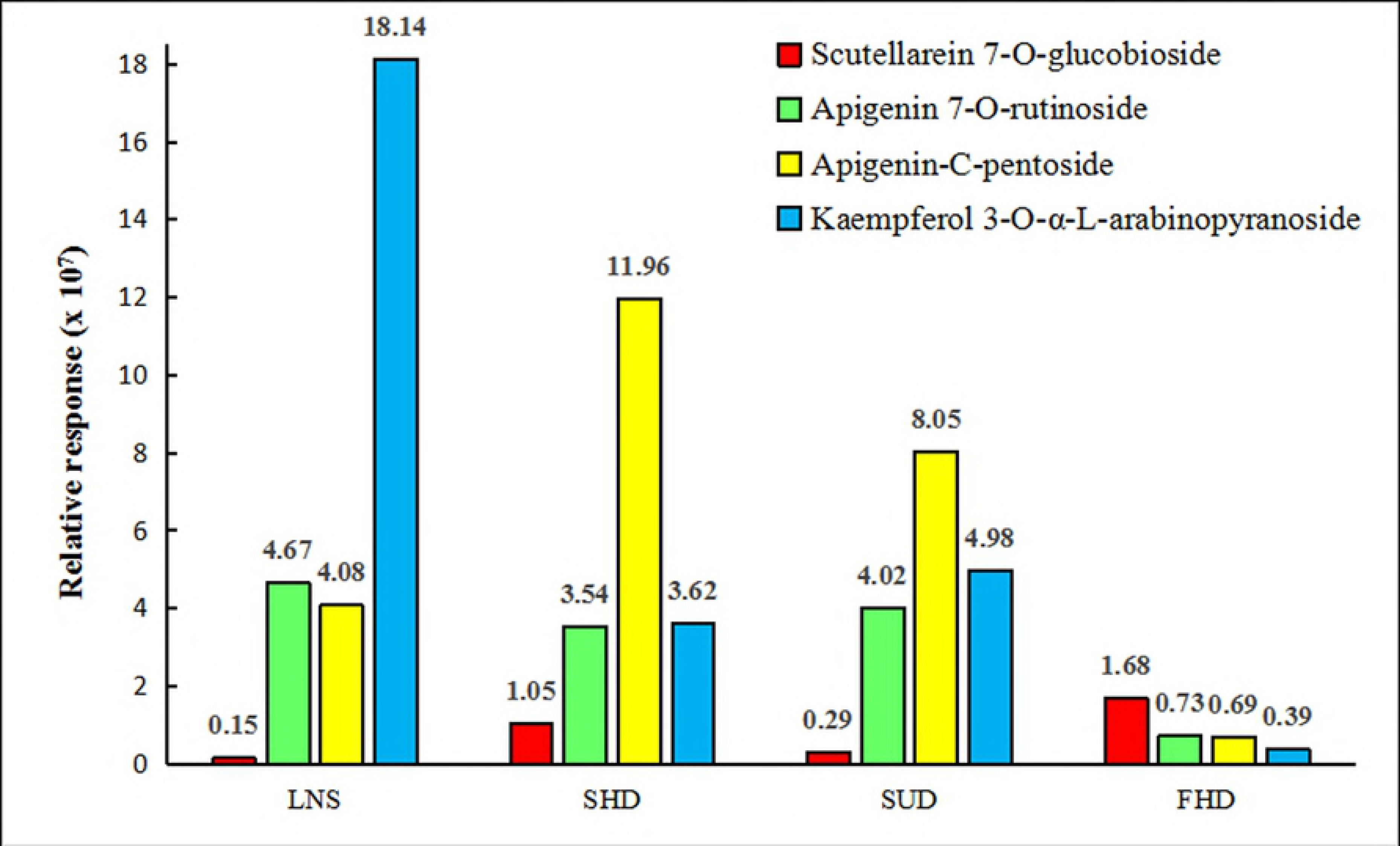

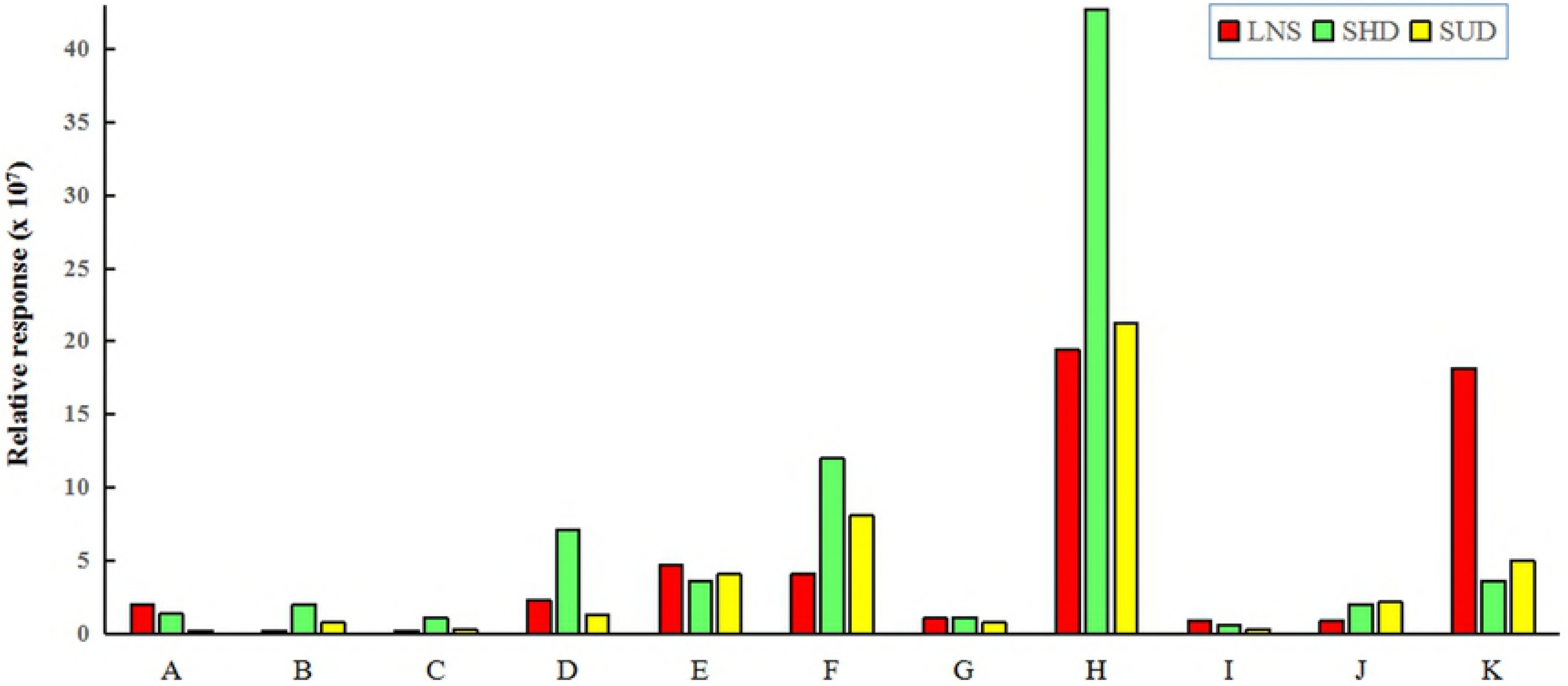

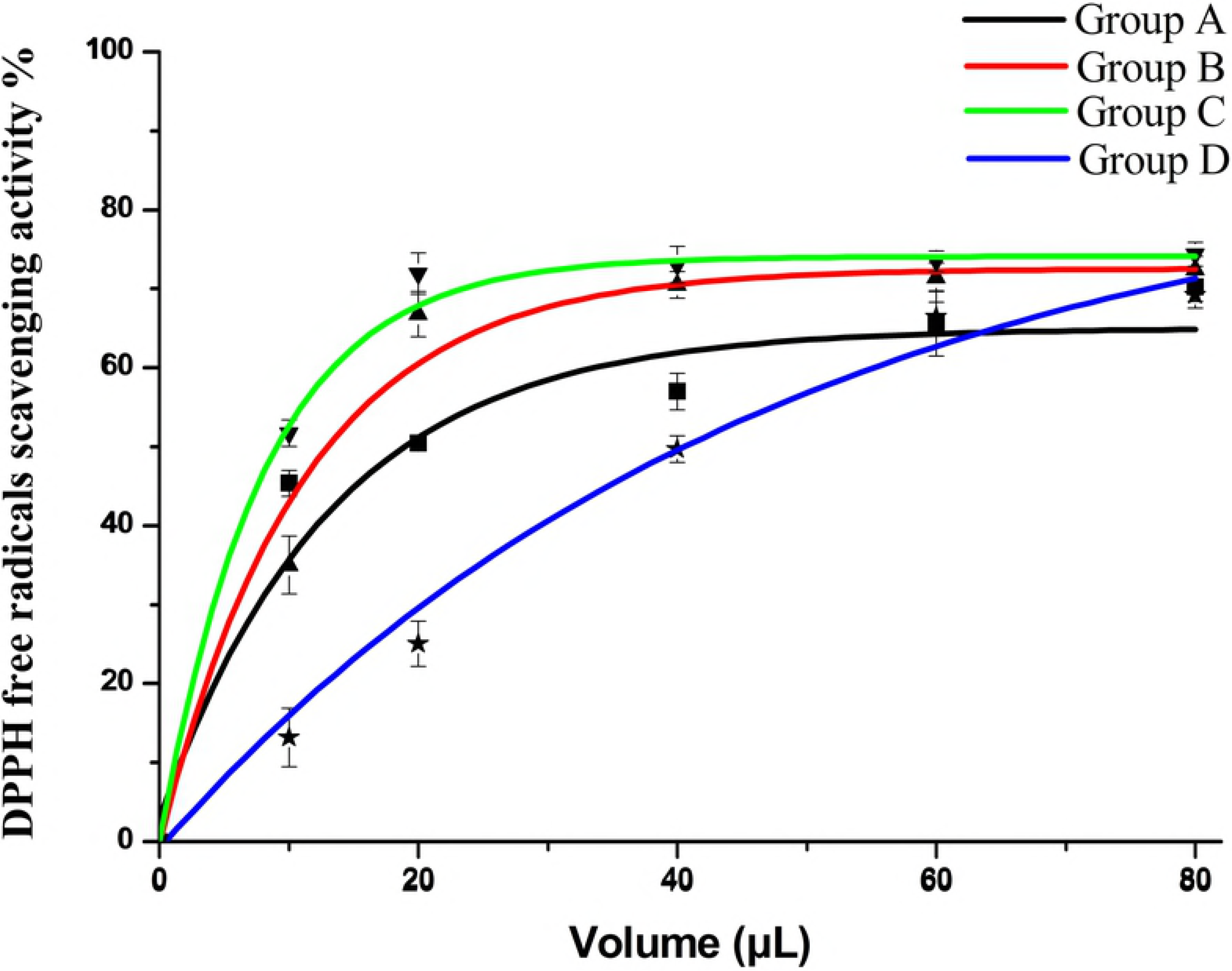

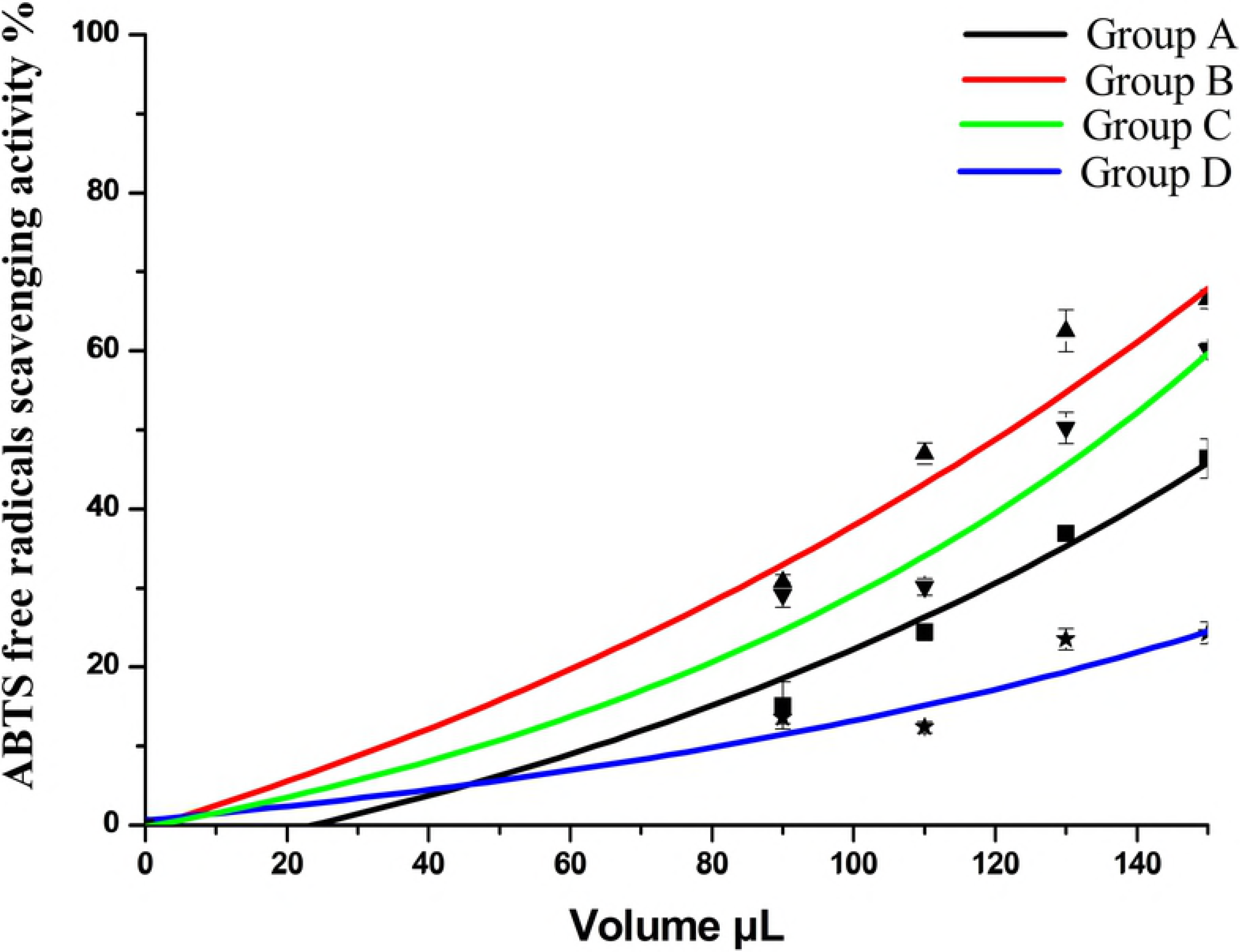

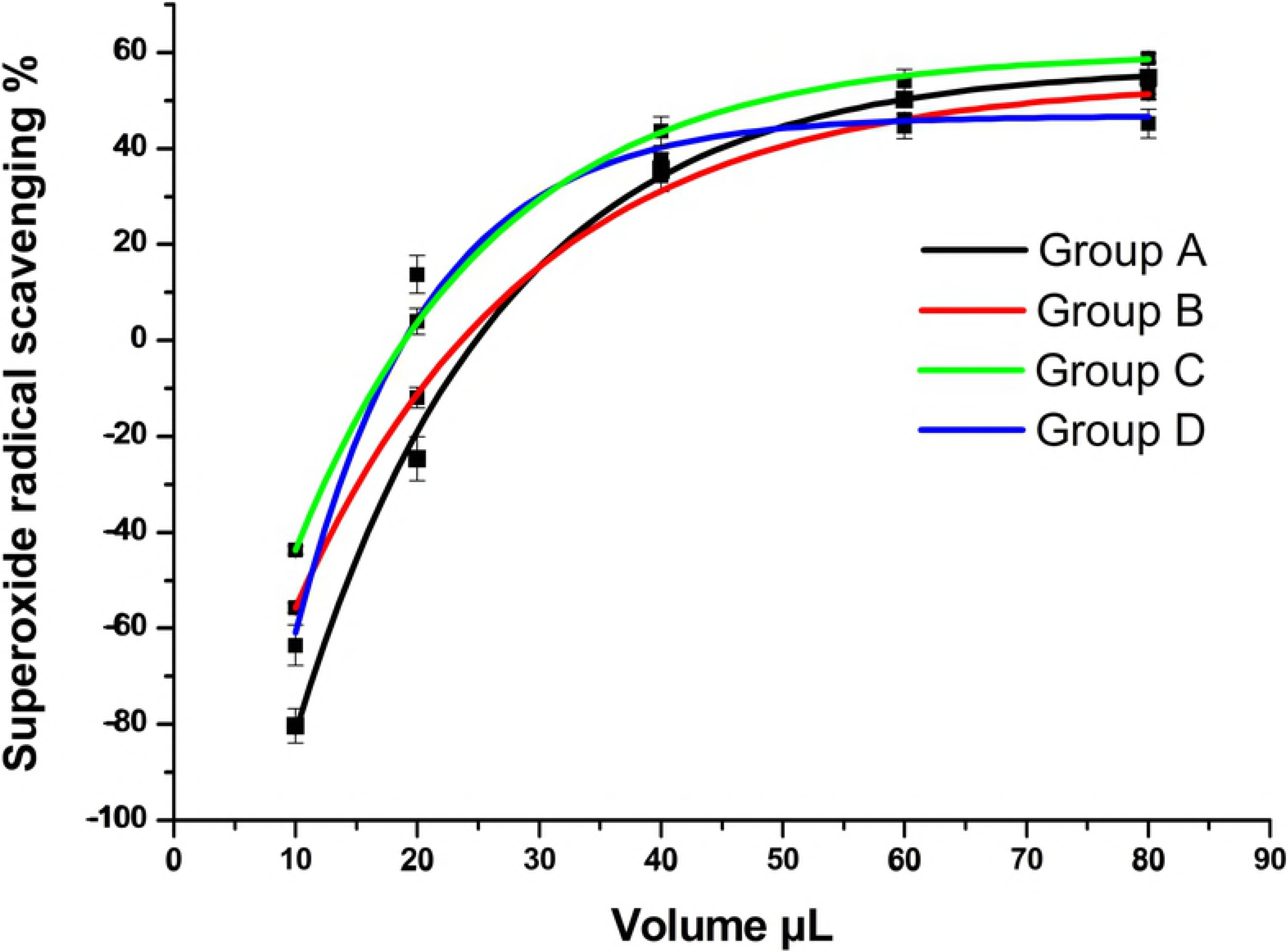

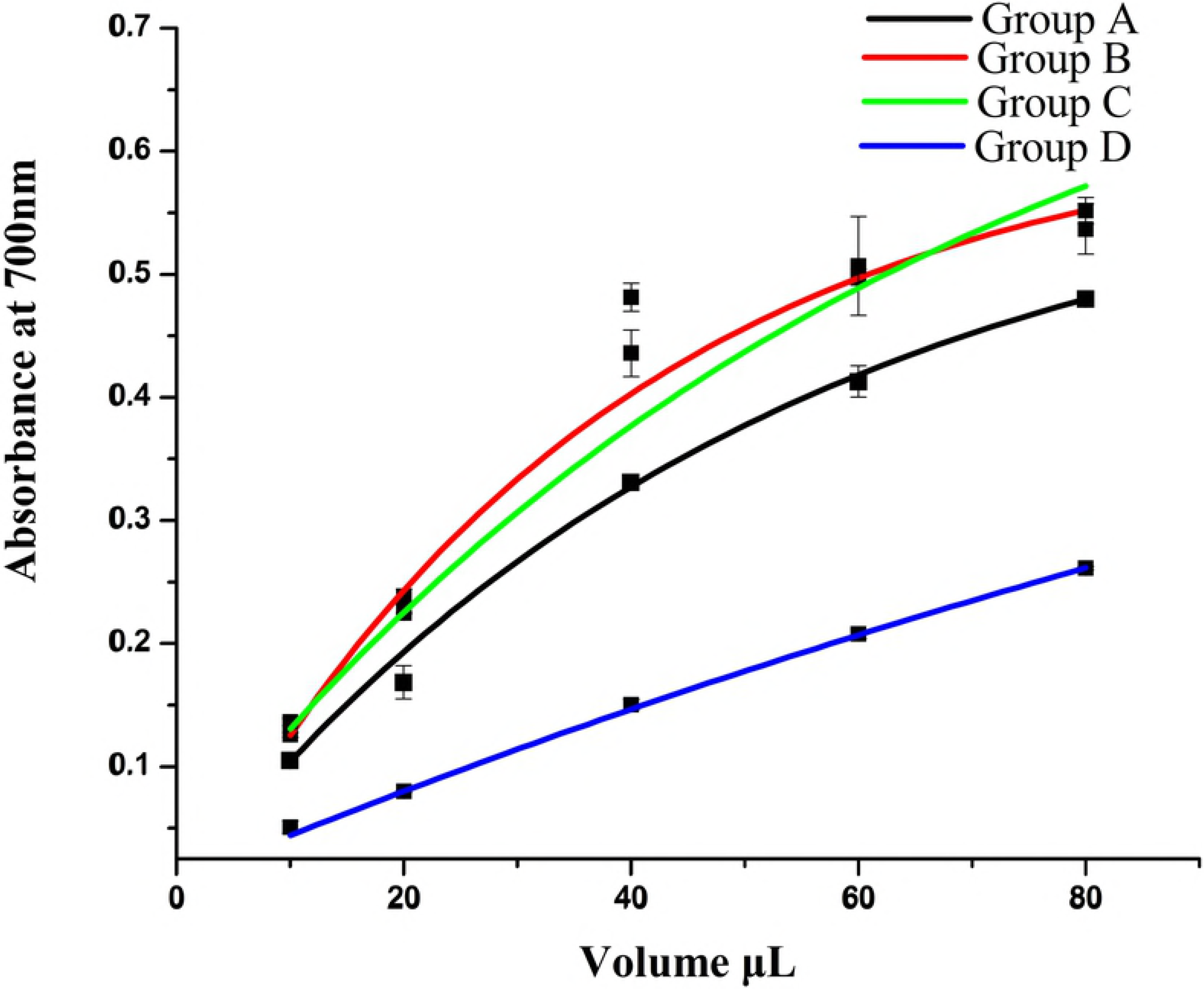

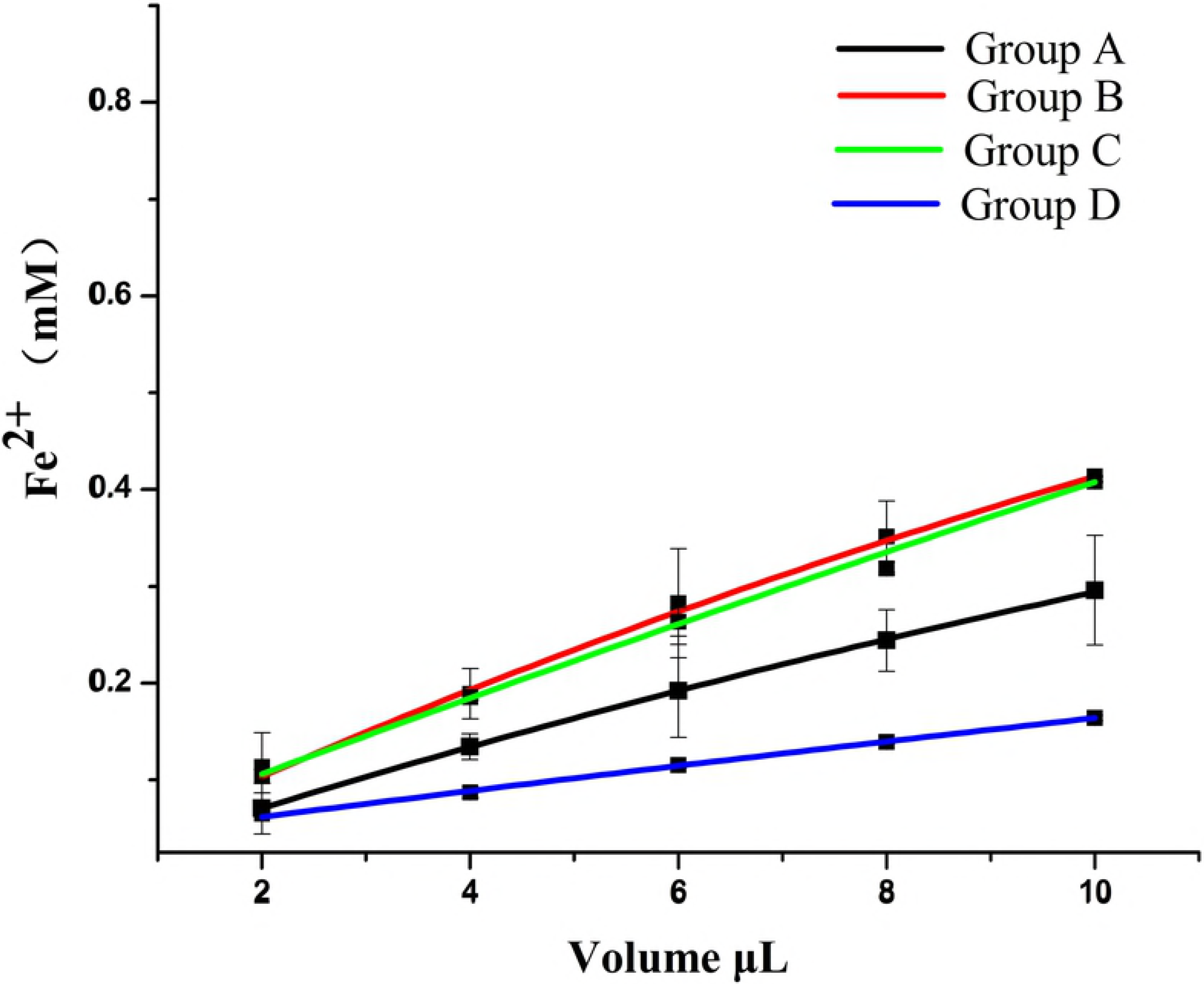

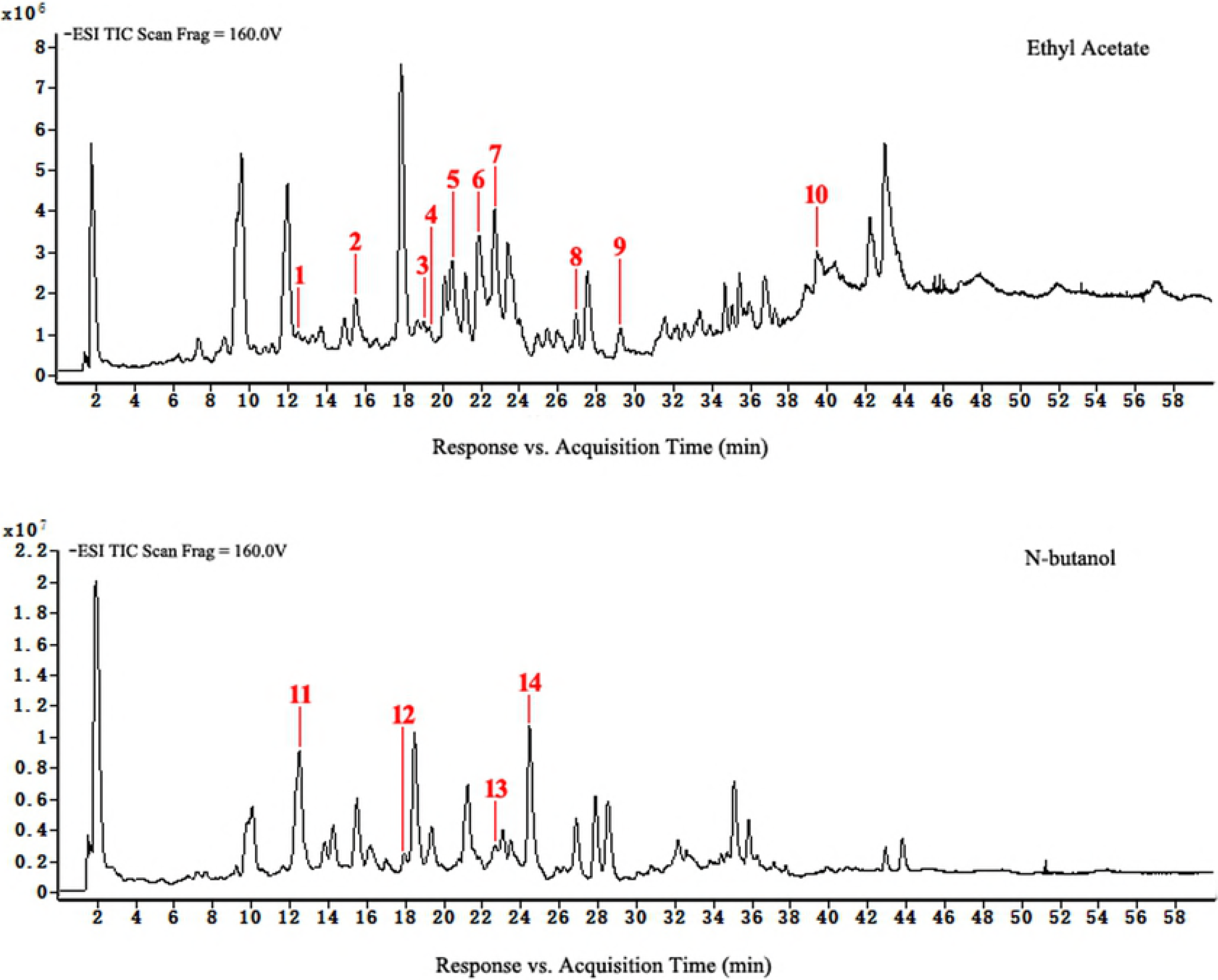

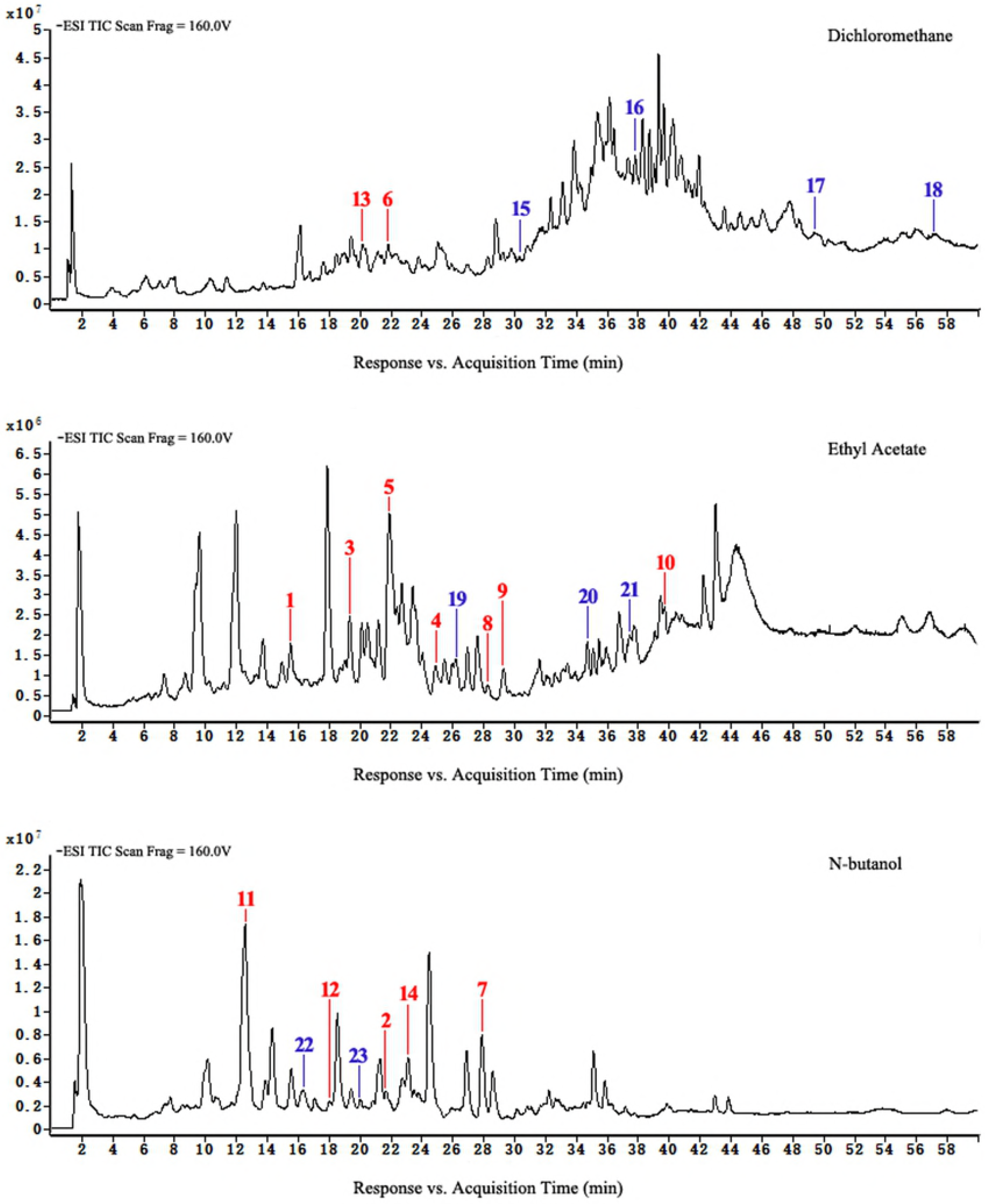

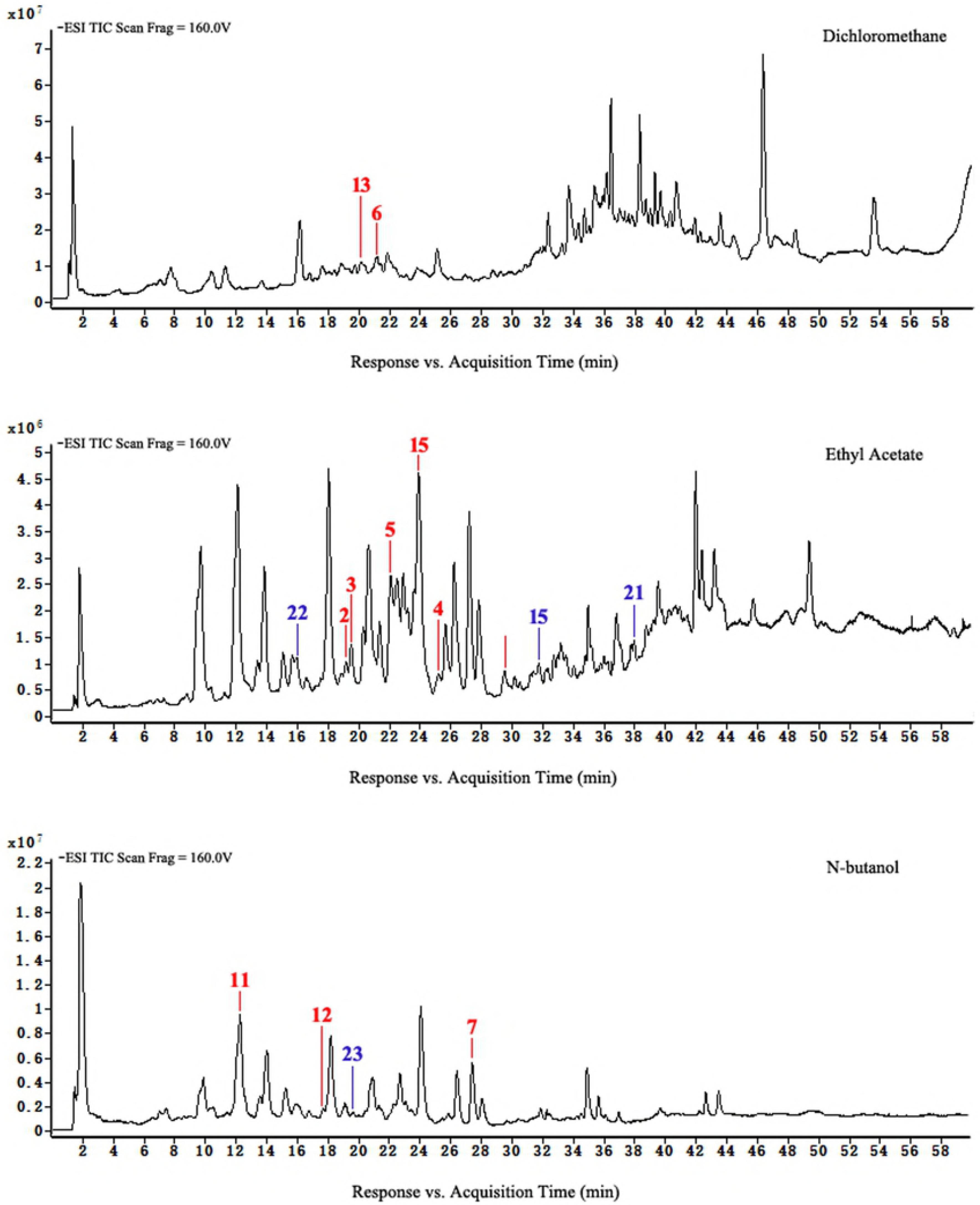

